# Aberration modeling in deep learning for volumetric reconstruction of light-field microscopy

**DOI:** 10.1101/2023.02.22.529610

**Authors:** You Zhou, Zhouyu Jin, Qianhui Zhao, Bo Xiong, Xun Cao

## Abstract

Optical aberration is a crucial issue in optical microscopes, which fundamentally limits the practical imaging performance. As a commonly encountered one, spherical aberration is introduced by the refractive index mismatches between samples and environments, which will cause problems like low contrast, blurring, and distortion in imaging. Light-field microscopy (LFM) has recently emerged as a powerful tool for fast volumetric imaging. The appearance of spherical aberration in LFM will cause large changes of the point spread function (PSF) and thus greatly affects the imaging performance. Here, we propose the aberration-modeling view-channel-depth (AM-VCD) network for LFM reconstruction, which can well mitigate the influence of large spherical aberration. By quantitatively estimating the spherical aberration in advance and modeling it in the network training, the AM-VCD can obtain aberration-corrected high-speed visualization of three-dimensional (3D) processes with uniform spatial resolution and real-time reconstruction speed. Without any hardware modification, our method provides a convenient way to directly observe the 3D dynamics of samples in solution. We demonstrate the capability of AM-VCD under a large refractive index mismatch with volumetric imaging of a large-scale fishbone of largemouth bass. We further investigate the capability of AM-VCD in real-time volumetric imaging of dynamic zebrafish for tracking neutrophil migration.

## 1 Introduction

Light-field microscopy (LFM) has recently served as a vital candidate for high-speed, large-scale, and long-time volumetric imaging for live samples, especially in observing morphological and functional dynamics such as neuronal activity^1, 2^, embryo development^3, 4^, and vascular transport^5^. Compared with other three-dimensional (3D) imaging methods recording a sequence of two-dimensional (2D) images to form a 3D volume such as light sheet microscopy (LSM)^6–8^, confocal microscopy^9, 10^, and two-photon microscopy (2PM)^2^. LFM benefits from the advantages of single-shot acquisition manner, easy-to-build optical set-up, and low phototoxicity^1–4, 11–13^. LFM utilizes a microlens array (MLA) inserted in the detection path of a commonly-used epi-fluorescence microscopy to encode the 3D volumetric information in a snapshot acquisition^13^. Then by postprocessing the snapshot with deconvolution^14^ or learning-based^15, 16^ methods, high-speed volumetric imaging of various applications can be obtained with promising performance. Among these methods, the view-channel-depth (VCD) network^16^ is recently proposed for artifact-removal, resolution-enhanced, and real-time reconstruction. However, although the LFM applies the sub-aperture acquisition of optical information by using the MLA, its reconstruction performance is still heavily affected by the imaging aberrations, especially the spherical aberration caused by the refractive index mismatch between samples, sample-immersed solutions, sample containers, and objective-lens-immersed mediums, which is typically unavoidable in high NA microscopic imaging systems^17, 18^. It is because the appearance of spherical aberration in LFM will cause large changes of the point spread function (PSF) while both the deconvolution and deep learning methods are sensitive to the accurate modeling of PSFs.

Many pieces of research have been proposed to solve the problem of aberrations in microscopy, such as the different adaptive optics (AO) methods applied in LSM, confocal, and 2PM^18–22^, which commonly utilize spatial light modulators (SLMs) or deformable mirrors (DMs) to modulate the light wavefront. Some works even model the aberration in deep learning networks^23–25^ to well ease its influence and improve the imaging quality. However, the aberration problem is rarely explored in the field of LFM^5, 26^. Wu et.al^5^ combinedly utilize a scanning LFM system and iterative digital adaptive optics (DAO) to realize subcellular resolution and long-term observation of 3D dynamics. In addition, Zhang et.al^26^ propose introducing spherical aberration to expand the depth of field (DOF) of scanning LFM and reduce artifacts near the original focal plane. But to the best of our knowledge, no research has been proposed to achieve the snapshot 3D observation of live dynamic samples at micrometer resolution under the condition of heavy spherical aberration.

In this work, we introduce the aberration modeling (mainly the spherical aberration) into the learning-based reconstruction process of LFM (VCD network^16^) to enhance imaging performance. By further integrating with automatic aberration evaluation and loss function optimization, our method, termed aberration-modeling view-channel-depth (AM-VCD) network, achieves resolution-uniform, artifact-reduction, and real-time LFM reconstruction, even in a situation with large spherical aberration. Our method does not make any changes to the hardware of LFM. Through modeling spherical aberration in network training, it is convenient and widely applicable in different LFM systems. The spherical aberration, modeled on the basis of Zernike polynomials, can be automatically estimated by measuring the PSFs of beads or bead-like structures inside the scene to be observed, as shown in Fig. 1 (see Section 2.3 and Fig. S2 for details). We first demonstrate the principle and improvement of our method through experiments of fluorescent beads, which can obtain aberration-corrected high-speed volumetric imaging even under a large spherical aberration. We also conduct some experiments under situations with varying degrees of spherical aberration to further verify the effectiveness and practicability of our method, where direct deconvolution and learning-based methods both fail to achieve a tolerable imaging quality. We realize the high-quality 3D imaging of largemouth bass fishbone in a volume of 426×426×101 μm^3^ with a measured spherical aberration of −0.9λ (20×/0.5NA air-immersion objective). We track the neutrophil migrations in dynamic zebrafishes in a volume of 426×426×101 μm^3^ with a measured spherical aberration of −0.5λ (20×/0.5NA air-immersion objective) with 30 ms exposure time and 6 Hz camera frame rate (which is limited by the hardware of the camera).

**Fig. 1.**
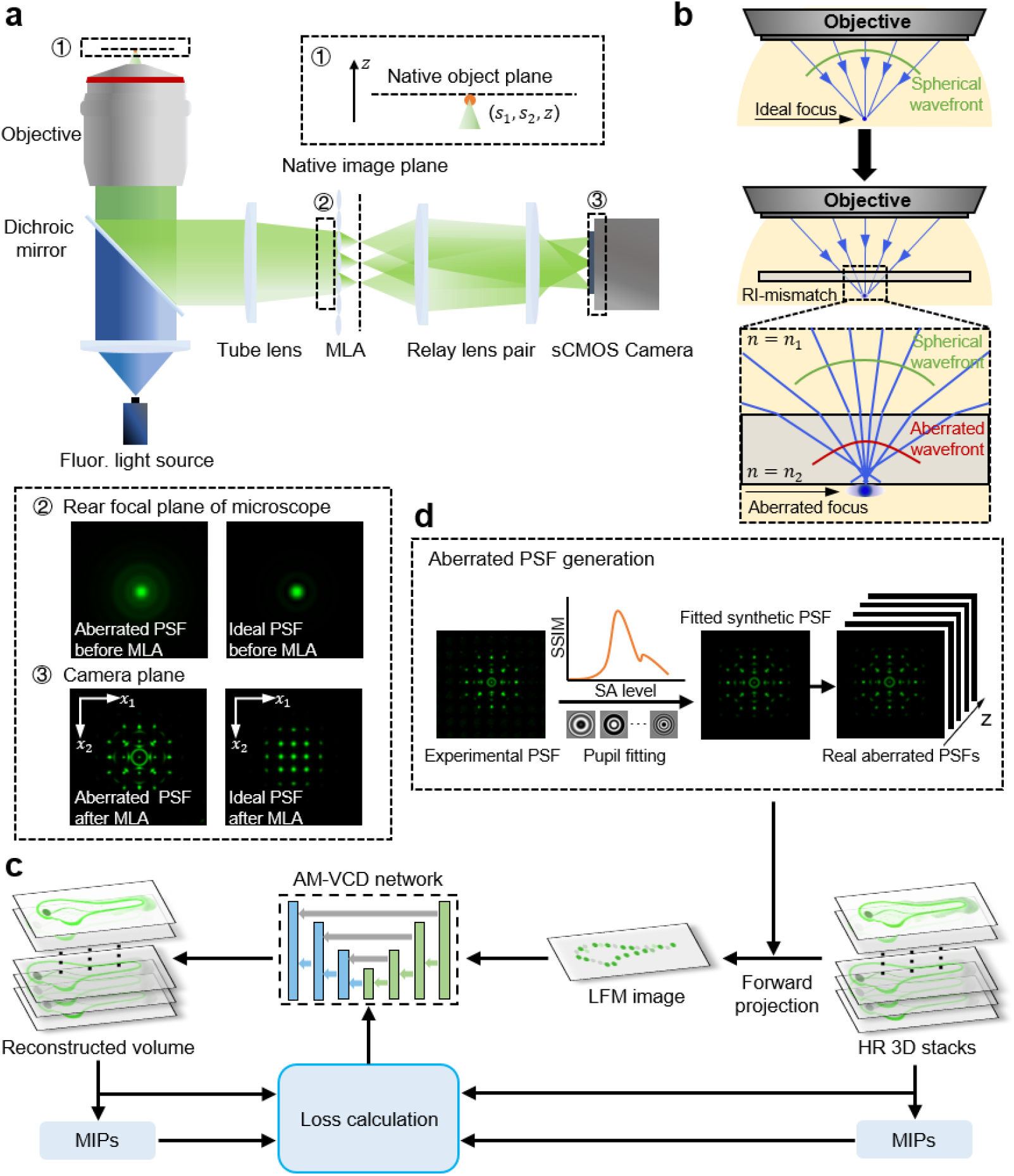
Principle of AM-VCD method. **a.** The optical set-up of light-field microscopy (LFM). A microlens array (MLA) is inserted at the rear focal plane of a conventional benchtop microscope to subsample the image of volumes captured by the front microscope system. The zoom-in panel 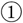 shows the relationship of a sample space point with the pattern in the native objective plane. The zoom-in panels 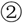 and 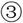 show the ideal (diffraction-limited) point spread function (PSF) and the aberrated PSF (i.e., spherical aberration) before and after the MLA, respectively. **b.** The spherical aberration is introduced by refractive index mismatches, which causes the failure of light to converge to a spot. **c.** The schematic diagram of how the proposed method eliminates the spherical aberration in LFM. Training pairs are generated from high-resolution (HR) 3D stacks with aberrated PSFs. The AM-VCD network is trained by iteratively minimizing the difference between reconstructed volumes and groundtruth volumes. The weighted difference between axial maximum intensity projections (MIPs) of reconstructed volumes and ground truths is also added in the loss function to ease reconstruction artifacts. **d.** The real aberrated PSFs are obtained by measuring the PSFs of fluorescence beads or bead-like structures in samples (see Fig. S2 for details).

## 2 Principles and Methods

### 2.1 Optical set-up

We build up a conventional LFM system in this work, which is appended to an epi-fluorescence microscope (Zeiss, Axio Observer 7) equipped with a fluorescence lamp illuminator (X-Cite 120Q) and a fluorescence filter set (Zeiss, Excitation: BP 450-490, Emission: LP 515). The optical set-up is shown in Fig. 1a. A 1:1 relay system is used to conjugate the back focal plane of MLA (RPC Photonics, MLA-S100-f21) with the detection plane of the camera sensor (PCO. edge 26 sCMOS camera, 5120×5120 pixels, and 2.5 μm pixel size). Each microlens covers around 11 × 11 effective pixels, which corresponds to the angular resolution. We bin 4×4 pixels of the camera sensor for image acquisition, resulting in a 10 μm effective pixel size. We use a 20× 0.5NA air-immersion objective lens (Zeiss Objective EC Plan-Neofluar 20×/0.50 M27) and a 63×/1.25NA oil-immersion objective lens (Zeiss Objective EC PN 63 ×/1.25 Oil M27) in experiments.

### 2.2 AM-VCD principle

The wave optical model of LFM proposed by Broxton et al.^14^ utilizes Scalar Debye Theory^27^ to describe the wavefront at the native image plane generated from a point source at the object space. This theory is based on the assumption that the PSF model is diffraction-limited and circularly symmetrical. However, when the system’s aberrations cannot be disregarded, the original PSF model becomes insufficient to accurately describe the real LFM system, which will heavily degrade the reconstruction quality of both deconvolution and learning-based methods. Therefore, we introduce the aberration modeling into the PSF generation of LFM and integrate it with the deep-learning method to achieve resolution-uniform, artifact-reduction, and real-time reconstruction. The proposed AM-VCD method can be mainly decomposed into two parts: aberration modeling and network reconstruction. In the following sections, we first mathematically represent the optical modeling of a conventional LFM and how we introduce aberration terms into this model. Then we describe the network realization of the proposed AM-VCD and how we evaluate the aberration in real experiments.

#### Forward imaging process of LFM

A conventional LFM system includes two optical hardware components: a benchtop wide-field fluorescent microscope and an MLA, as illustrated in Fig. 1a. We designate the object space coordinates as (*s*_1_, *s*_2_, *z*) and the sensor plane coordinates as (*x*_1_, *x*_2_). The original PSF of LFM at the sensor plane can be expressed as

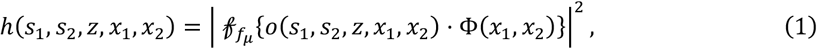

where 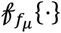 represents the Fresnel diffraction operator propagating a distance *f_μ_* along the optical axis. *o*(*s*_1_, *s*_2_, *z,x*_1_, *x*_2_) is the optical field at the native image plane (NIP) generated by a point source *s*(*s*_1_, *s*_2_, *z*) in the object space, which can be formulated by the Scalar Debye Theory as

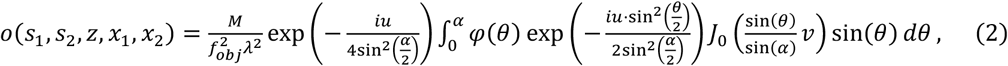

where *M*is the magnification of the objective, *f_obj_* is the focal length of the objective, *λ* is the emission wavelength, *α* is the maximum angle corresponding to NA of the objective. *φ*(*θ*) indicates the apodization function of the microscope, *i* is the imaginary sign, and *J*_0_ represents the zeroth order Bessel function of the first kind. 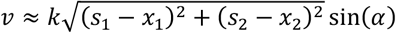 and 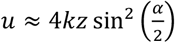 are normalized radial and axial optical coordinates, where 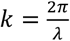 is the wave number.

Besides, in Equation (1), Φ(*x*_1_, *x*_2_) represents the phase modulation mask of MLA, whose pitch size is *d* and focal length is *f_μ_*, and can be expressed as

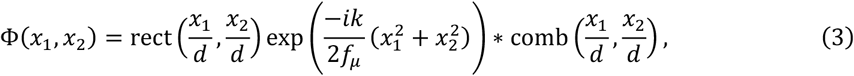

where rect(·) indicates rectangle function, comb(·) indicates 2D comb function, and * denotes 2D convolution.

#### Aberration modeling in LFM

To introduce aberration terms into the LFM imaging model, we apply phase modulation at the pupil plane^5, 21^ rather than modifying the formula of Scalar Debye Theory for its efficiency in computation, generality in introducing different types of aberrations, and convenience in integrating the Zernike polynomials.

The pupil plane coordinates are denoted as (*ρ, θ*). We use the forward and inverse Fourier transform to introduce a virtual 4-f system into the LFM system, which enables us to do phase modulation at the system’s pupil plane. Then we use the Zernike polynomials to fit the aberration, which is commonly used to describe the optical path difference (OPD) of the wavefront at the pupil plane^28^. The modulated or aberrated pupil function 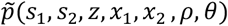 can be expressed as follows

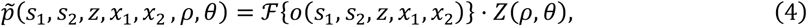

where 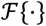 denotes the 2D Fourier transform and *Z*(*ρ, θ*) is the aberrated wavefront, which can be represented by the Zernike polynomials. Then the aberrated optical field 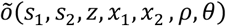 at the NIP can be calculated as

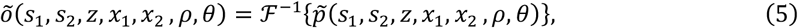

where 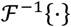 denotes the inverse 2D Fourier transform. Therefore, the aberrated PSF of LFM at the sensor plane can be expressed as

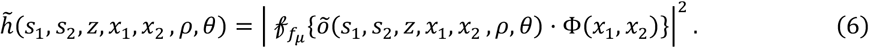

#### AM-VCD network realization

As mentioned, the VCD network^16^ is a recently proposed deep learning method for real-time and artifact-removal reconstruction of LFM. We integrate the aberration modeling of LFM into the original VCD network to realize the AM-VCD network. Specifically, as shown in Fig. 1c-d, we first calculate PSFs of the aberrated LFM according to Equation (6), where the system parameters are the same as those in experiments including the real aberrations. Similar to the VCD network, we then acquire high-resolution 3D volumes by confocal microscopy or by simulation synthesis as the ground truths and subsequently do 3D convolutions to the ground-truth 3D volumes with the aberrated PSFs (i.e., the forward imaging model of LFM) to generate corresponding aberrated LFM images. The paired ground-truth 3D volumes and aberrated LFM images are applied for our AM-VCD network training.

Resolving the 3D structure of an object from a single-shot LFM image is an ill-posed problem and the loss function of the network will highly affect the reconstruction performance. We thus modify the loss function of the original VCD network to ease the remained artifacts and balance the energy distribution axially, especially when the sample distribution is dense and complicated. Specifically, the VCD network uses the mean square error (MSE) between VCD inferences and ground-truth 3D volumes as the loss function^16^, which can be expressed as

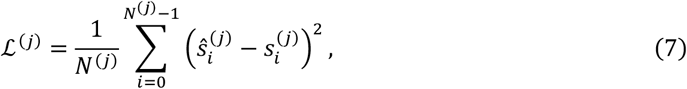

where 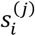 represents the *i*-th voxel of the ground truth in the *j*-th pair of the training set and 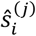 represents the *i*-th voxel of the reconstructed 3D distribution in the *j*-tli pair of the training set. *N*^(*j*)^ indicates the total voxel numbers of the 3D Volume ***s***^(*j*)^.

Apart from the MSE between the paired volumes, we also impose a secondary constraint based on the axial information of the 3D object during the training of the network. We calculate the maximum intensity projections (MIPs) in the x-z, y-z, and x-y directions of the reconstructed 3D distribution and compare the differences between these MIPs and their corresponding ground truths. We add this MIP constraint to the loss function as follows

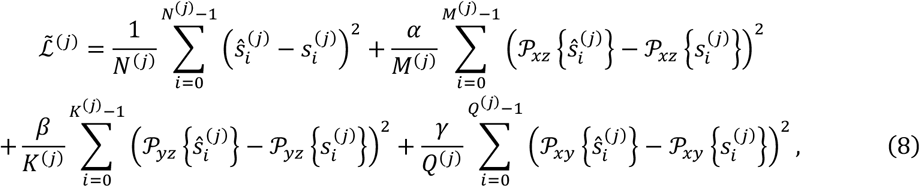

where 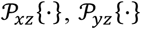, and 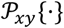 represent the x-z, y-z, and x-y direction MIPs of the 3D object, respectively. *α*, *β*, and *γ* are three pre-set hyperparameters as weights, which we set *α* = *β* = 1 and *γ* = 0 here. *N*^(*j*)^ is the total voxel numbers of 3D volume ***s***^(*j*)^. *M*^(*j*)^, *K*^(*j*)^, and *Q*^(*j*)^ represent total voxel numbers of 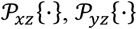, and 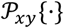, respectively.

### 2.3 Aberration evaluation

Evaluating the real aberrations in experiments is a crucial task in the aberration-corrupted LFM system, which will determine the accuracy of PSF modeling and thus affect the final reconstruction performance. The main steps of aberration evaluation of our method are as shown in Fig. 1d. We first add different levels of aberrations into the PSFs of the original LFM, generating a series of aberrated PSFs according to Equation (6), where the parameters are set the same as those in real experiments. In this work, we only consider the most commonly encountered spherical aberration, which is introduced by the refractive index mismatch between samples and imaging environments. Given a certain level of spherical aberration, its aberrated pupil function can be directly generated by the Zernike polynomials (see Supplementary Note 1 for details).

Then we obtain an experimental PSF image by taking a photograph of a fluorescent bead arbitrarily immersed in the sample medium or a bead-like structure inside the sample, which has a smaller than the resolution limit of LFM (e.g., 4-μm diameter in 20× experiments and 1-μm diameter in 63× experiments). We generate a series of synthetic PSFs with varying spherical aberration levels within an estimated depth. As shown in Fig. 1d, we then compare the synthetic PSFs with the captured experimental PSF image to estimate the level of the real spherical aberration (see Fig. S2 for details). Here we calculate the structural similarity index (SSIM) between each synthetic PSF and the experimental PSF to evaluate the real aberration as follows

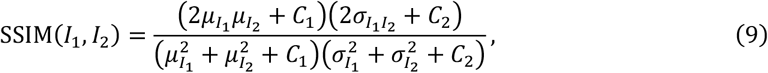

where *I*_1_ and *I*_2_ are normalized grayscale images, *μ*_*I*_1__ and *μ*_*I*_2__ are the local means of *I_1_* and *I_2_*, 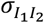 is the covariance of images *I_1_* and *I*_2_, 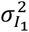 and 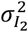 are variances of *I_1_* and *I*_2_, and *C*_1_ and *C_2_* are constants to avoid a division by null. The real spherical aberration in experiments is corresponding to the maximum SSIM value, which is further used to generate all the PSFs of the aberrated LFM system according to Equation (6) (see Fig. S2 and Fig. S3 for details).

## 3 Results

### 3.1 Resolution evaluation of AM-VCD by imaging fluorescent beads

We first test the imaging performance, especially the obtainable spatial resolution both laterally and axially, of our method by imaging the fluorescent beads. We apply an oil-immersion objective lens (Zeiss Objective EC PN 63 ×/1.25 Oil M27) to image 1-μm diameter fluorescent beads (ABT-18-3-01, Bitoyscience), which are placed in a glass bottom dish (15-mm diameter and 0.17-mm bottom thickness, NEST). The space between the objective lens and the dish is immersed with silicone oil. According to Section 2.3, we can estimate the level of the real spherical aberration (0.3λ here) in this experiment in advance. Noting that this spherical aberration inherently exists in the optical system and will heavily affect the reconstruction quality of LFM, which is caused by the refractive index mismatch between the objective-immersed silicone oil (refractive index = 1.518) and the disk bottom well (refractive index = 1.523).

#### Synthetic dataset generation for network training

We synthesize the datasets of fluorescent beads to train both the VCD and our AM-VCD networks. We first simulate many binary beads with isotropic resolutions and make them randomly distributed in a 3D space within a 30-μm depth range. We then apply a 3D-Gaussian kernel to convolve these beads to finely control their full width at half maximum (FWHM) values to be 1× 1× 1 μm^3^. By this, we obtain a 3D stack of randomly distributed beads, which is regarded as the ground-truth volume. We totally generate about 4000 3D stacks to form the groundtruth dataset. Next, we calculate the PSFs of the original LFM according to Equation (1) for VCD and PSFs of the aberrated LFM according to Equation (6) for AM-VCD. We subsequently convolve the ground-truth 3D stacks with original PSFs and aberrated PSFs respectively to acquire the corresponding LFM images for VCD and AM-VCD. Finally, these two paired datasets are respectively applied to train the VCD and AM-VCD networks.

#### Experiment evaluation

We prepare a 3D distributed sample of fluorescent beads and experimentally capture an LFM image of the sample. We reconstruct the 3D image stacks of beads by using Richardson-Lucy (RL) deconvolution^14^, VCD^16^, and our AM-VCD. We also integrate our aberration model into RL deconvolution, termed AM-RL deconvolution, to reconstruct the beads for better comparison. As shown in Fig. 2a-d, aberration-modeling-based methods (both AM-RL deconvolution and AM-VCD) can improve reconstruction resolution when aberration exists. AM-RL deconvolution can effectively suppress artifacts that exist in the results of conventional RL deconvolution (Fig. 2a-b). AM-VCD network can further improve the spatial resolution both laterally and axially, compared to the VCD network and AM-RL deconvolution, and it suppresses the reconstruction errors in the result of the VCD network, as marked by yellow arrows in Fig. 2c. Noting that in Fig. 2e, the reconstructed beads by the RL deconvolution and VCD network have some axial shifts compared to the reconstruction results by aberration-modeling methods. It indicates that reconstruction without considering spherical aberration will lead to incorrect axial positioning of sample structures, which is consistent with the characteristic of spherical aberration. Our AM-VCD network achieves an isotropic resolution of about 1.4 μm here.

**Fig. 2.**
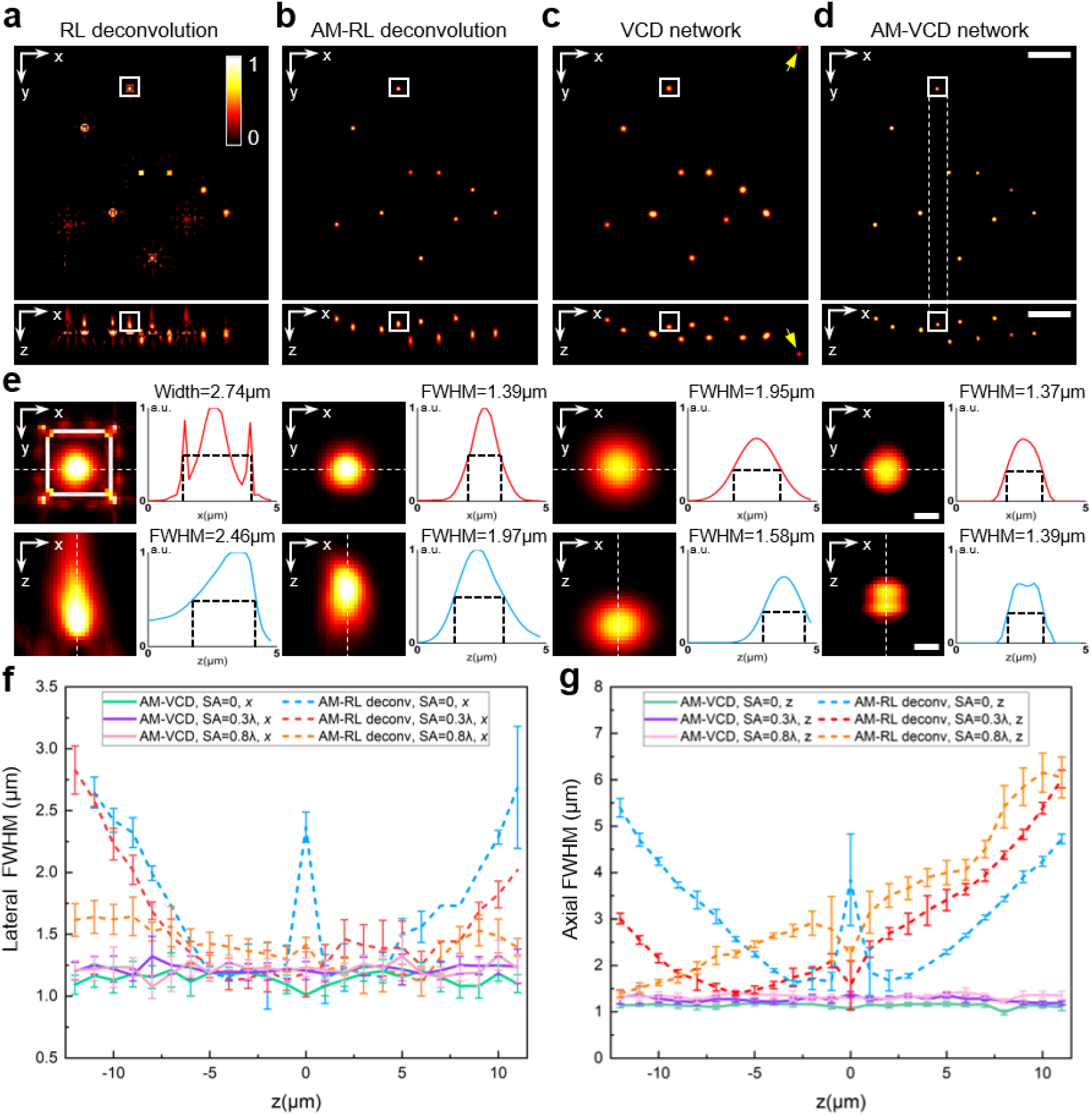
Performance and resolution evaluation of AM-VCD network by imaging fluorescent beads with an oilimmersion objective lens (63×/1.25NA). **a-d.** Lateral and axial MIPs of real-captured fluorescent beads reconstructed by RL deconvolution, AM-RL deconvolution, VCD network, and AM-VCD network, respectively. **e**. Zoom-in areas marked by white boxes in **a-d**. The profiles along dashed lines are plotted and full width at half maximum (FWHM) values are calculated in both lateral and axial directions. a.u., arbitrary unit. **f-g.** Average lateral and axial FWHM curves of the simulated 3D-distributed beads under different spherical aberration (SA) levels (0, 0.3λ, and 0.8λ) are plotted, which are reconstructed by AM-RL deconvolutions (dashed lines) and AM-VCD network (solid lines) using the corresponding aberrated PSFs. In both experiments and simulations, aberration-modeling-based methods (both AM-RL deconvolution and AM-VCD network) improve the reconstruction resolutions. AM-VCD network can further improve the spatial resolution both laterally and axially and can suppress the reconstruction errors that occurred in the results of the original VCD network. Scale bar, 10 μm **a-d**, 1 μm **e**.

#### Simulation evaluation

We also use simulations of beads under different levels of spherical aberration (SA = 0, 0.3λ, and 0.8λ) to evaluate the effectiveness of our method. The test LFM images are synthesized by randomly placing 100 beads at different layers of a 3D stack and then applying the LFM forward model to process the 3D stack using corresponding aberrated PSFs. Since conventional RL deconvolution and VCD network will introduce some artifacts and axial shifts in reconstruction as having been demonstrated in Fig. 2a, 2c, and 2e, their FWHMs can hardly reflect the actual imaging resolution. Thus in simulations, we only test the resolution-resolved ability of AM-RL deconvolution and AM-VCD network. For training the AM-VCD network, we generate paired ground-truth 3D stacks and LFM images using the above-mentioned synthetic dataset generation steps with different aberrated PSFs. As the reconstructed results shown in Fig. 2f and 2g, we can find that for systems with the positive spherical aberration, the depth of the highest lateral resolution achieved by AM-RL deconvolution moves away from the focal plane in a certain direction as the spherical aberration increases and the region near the focal plane is no longer with the lowest reconstruction resolution (see also Fig. S10). We think it is due to the change of the spatial sampling patterns in different depths, caused by wavefront modulation of the spherical aberration. In comparison, the AM-VCD network obtains relatively uniform and almost isotropic resolution across over 25-μm imaging depth under different levels of spherical aberration. It reaches an average resolution of 1.1 μm (*x, y*) and 1.1 μm (z) when SA = 0, 1.2 μm (x,y) and 1.2 μm (z) when SA = 0.3λ, and 1.2 μm (x,y) and 1.3 μm (z) when SA = 0.8λ. The results show that the AM-VCD network has a stable performance even under a large spherical aberration level (SA=0.8λ) and is able to achieve a resolution close to the case without aberration (SA = 0).

Besides, we also conduct all the above experiments and simulations using an air-immersion objective lens (20×/0.5NA) to demonstrate the effectiveness and generality of our method, as the results shown in Fig. S4.

### 3.2 AM-VCD with MIP constraints can further suppress periodic artifacts

In this experiment, we compare different reconstruction performances by RL deconvolution, AM-RL deconvolution, VCD, and AM-VCD in imaging biological samples with complex structures. Here we train the AM-VCD network with and without MIP constraints respectively to demonstrate the validity of the loss function modification (adding MIP constraints) in Equation (8). It is noted that in the rest of the paper, the loss functions applied in AM-VCD training are all equipped with MIP constraints if there is no special explanation.

We first use a 63 ×/1.25NA oil-immersion objective to image the ring-like structures of dandelion villi slice (commercially bought from AOXING Laboratory Equipment). The space between the objective and the specimen is filled with water to artificially introduce a large spherical aberration, which is evaluated to be 0.8λ. The captured LFM image is shown in Fig. 3g. We place some fluorescent beads on the cover slip to capture the experimental PSF for aberration evaluation, which is 0.8λ after estimation. From the reconstructed results in Fig. 3a-f, we observe that methods such as the RL deconvolution, our modified AM-RL deconvolution, and the original VCD network can hardly reconstruct the 3D distribution of the sample due to the introduced large spherical aberration, especially in axial directions. AM-VCD network without MIP constraints performs much better in correctly resolving the sample structures and shows a much better imaging resolution referring to the ground truth captured by a confocal microscope (Zeiss, LSM 880). But it still produces some periodic artifacts and some errors in axial energy distribution, due to the object’s complex 3D structures. In comparison, the AM-VCD network trained with MIP constraints can further improve the imaging quality and well suppress the periodic artifacts and axial errors in the reconstructed results, demonstrating the validity of the loss function modification, as shown in Fig. 3d-e and 3i-j. It achieves the best reconstruction performance and highest spatial resolution, especially in the axial direction. The artifact-suppress performance of AM-VCD with MIP constraints can also be seen in the Fourier domain, as shown in Fig. 3h. In addition, for computational cost, the deconvolution methods need several minutes of reconstruction time (about 20 minutes for a 671 × 671 LFM image here), while the network methods only need several tens of milliseconds to reconstruct a 3D volume (about 70 ms).

**Fig. 3.**
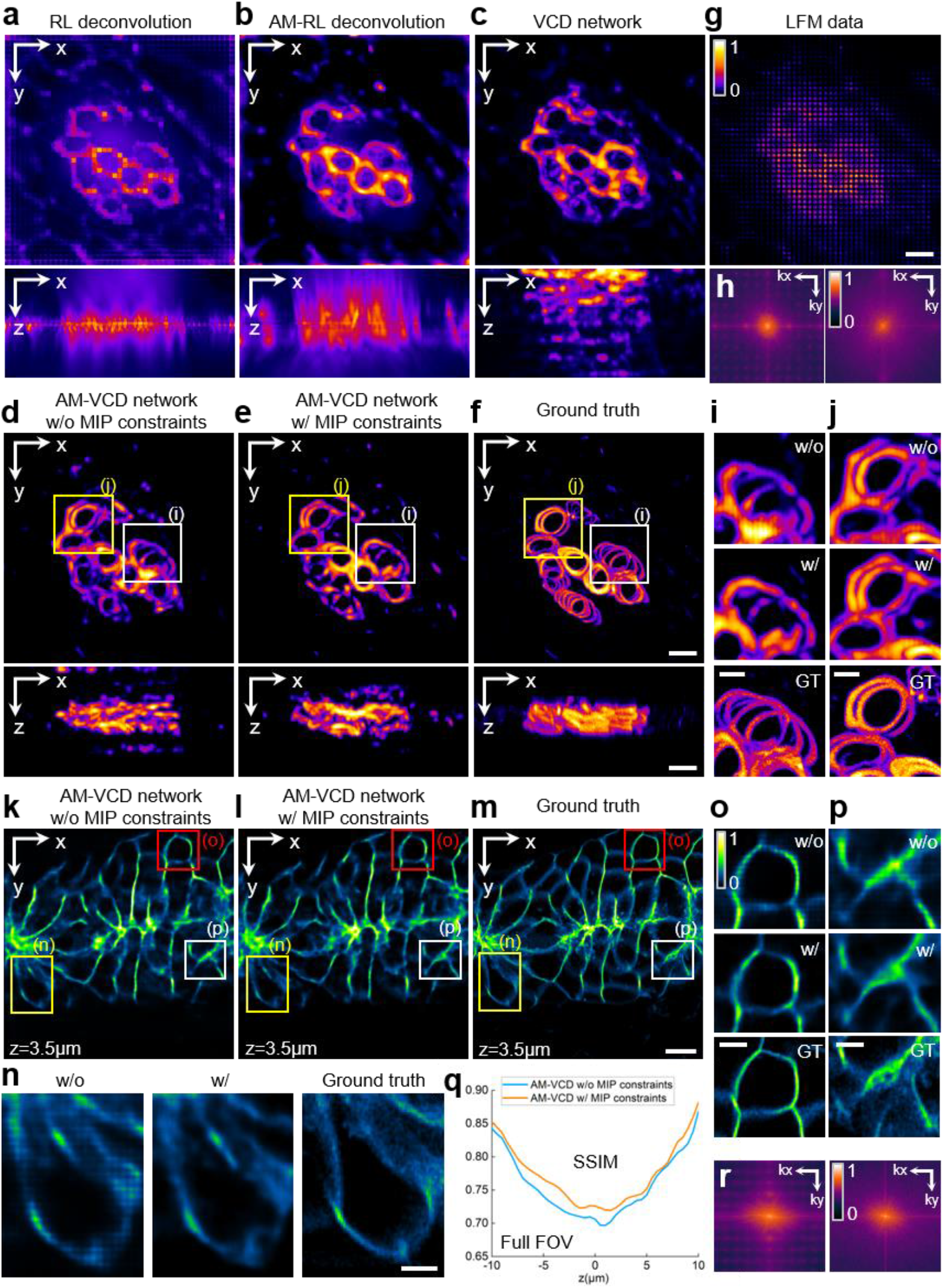
Performance validation of MIP constraints in AM-VCD by imaging biological samples with an oil-immersion objective lens (63×/1.25NA). **a-e.** MIPs of reconstructed dandelion villi slice with ring-like structures by RL deconvolution, AM-RL deconvolution, VCD, AM-VCD without MIP constraints, and AM-VCD with MIP constraints. **f.** The MIPs of the ground truth image captured by confocal (Zeiss, LSM 880). To exhibit the strong capability of our AM-VCD network, we use a 63 ×/1.25NA oil-immersion objective to image samples, but replace the oil with water to introduce a heavy refractive index mismatch (i.e., large spherical aberration) in the LFM acquisition. The estimated spherical aberration is about 0.8λ. **g.** The raw LFM image captured by our LFM system. **h.** Fourier transform of **d** and **e**. **i.** Zoom-in areas marked by the yellow boxes in **d-f**. **j.** Zoom-in areas marked by the white boxes in **d-f. k-m.** A certain reconstructed slice of synthetic membrane data^20^ at z = 3.5 μm by AM-VC without and with (w/o and w/) MIP constraints and the corresponding ground truth. **n-p.** Zoom-in areas in **k-m** marked by yellow, red, and white boxes, respectively. **q.** the structural similarity index (SSIM) of the full field of view (FOV) membrane data reconstructed by AM-VCD network w/o and w/ MIP constraints across the depth range, referring to the ground truth. **r.** Fourier transform of **k** and **l**. Our proposed AM-VCD network w/ MIP constraints can suppress periodic artifacts of reconstructed results, which are illustrated in both the spatial and Fourier domain images, and achieve high-quality reconstruction performance both laterally and axially. Scale bar, 15 μm **a-g**, 7.5 μm **i-j**, 10 μm **k-m**, and 3 μm **n-p**.

To further verify the artifact-suppress ability of AM-VCD with MIP constraints, we borrow the 3D cellular membrane volume data^29^ to synthesize the training dataset and testing data by adding a spherical aberration level of 0.8λ. Since the cellular membrane data have quite complex structures, some periodic artifacts are introduced in the reconstrued result by AM-VCD without MIP constraints, as the reconstrued slice image (z = 3.5 μm) shown in Fig. 3k and some zoom-in areas shown in Fig. 3n-p. The periodic artifacts can also be found in the Fourier transform of Fig. 3k, as shown in Fig. 3r. On the contrary, AM-VCD with MIP constraints can greatly suppress the periodic artifacts in the reconstruction result, showing more consistent performance with the ground truth, as the reconstructed slice shown in Fig. 3l and zoom-in areas shown in Fig. 3n-p. The artifact-suppress ability can also be found in the Fourier transform of Fig. 3k and 3l, as shown in Fig. 3r. The SSIM curves in Fig. 3q quantitatively exhibit the improvement by adding the MIP constraints. Inheriting from the original VCD network, AM-VCD without MIP constraints performs well in reconstructing simple dot-like or line-like structures of samples. But it will inevitably produce some periodic artifacts when the 3D distribution of the object has more complex structures (e.g., dense distributions or complex surfaces). We demonstrate here that these periodic artifacts can be effectively removed by adding the MIP constraints.

### 3.3 AM-VCD improves fidelity in observing specimen immersed in a solution

We further verify that our AM-VCD network can directly image samples immersed in a solution that introduces a large spherical aberration, with high reconstruction accuracy. Many biological samples, especially living samples, always need to be placed in a solution (such as agar and water) for observation. However, to directly image a sample immersed in a solution is typically a challenging issue, since a heavy refractive index mismatch will be introduced between the sample solution and the objective-immersed medium, resulting in the large spherical aberration. We can try to reduce the refractive index difference by using matched objective-immersed media, but this way is not always possible and may need complicated medications of the optical system. Besides, some solvents are corrosive.

Our proposed AM-VCD network can well solve this problem without the need for hardware modification to the LFM system. In Fig. 4, we show that the AM-VCD is suitable to image the semitransparent hard bone of glycerol-immersed (n = 1.4746) largemouth bass sample by conveniently using an air-immersion 20×/0.5NA objective lens (Zeiss Objective EC Plan-Neofluar 20×/0.50 M27) on a typical inverted microscope, getting rid of the complex and delicate system with a water-immersed objective. We cut a small piece from the specimen and place it in the original solution (i.e. the glycerol) for observation, as shown in Fig. 4a. We randomly drop some green fluorescent beads into the sample solution to measure the aberration level. The sample has complex surfaces and the captured raw LFM images are under large spherical aberration (the estimated parameter is −0.9λ), which makes it hard for conventional LFM algorithms to recover. But our proposed AM-VCD obtains quite similar performance to the ground truth obtained by confocal, where the hole structures of the fishbone are successfully recovered with high accuracy and few artifacts. The 3D rendering images with a 426×426×101 μm^3^ volume are shown in Fig. 4b. In comparison, deconvolution methods such as RL deconvolution and our modified AM-RL deconvolution are unable to accurately reconstruct the structures of the sample and their achievable resolutions are relatively lower. The results of the VCD network have weird 3D structures and introduce large artifacts (e.g., the areas marked by white arrows in Fig. 4b). Additionally, as the slice images shown in Fig. 4c, our AM-VCD network demonstrates its strong optical sectioning ability (e.g., the areas marked by blue and yellow arrows in Fig. 4c). The reconstructed slice images by our method are well consistent with the ground truths in corresponding depths (z = −36 μm, 0 μm, and 36 μm), compared with other methods. This experiment verifies that our AM-VCD improves the fidelity in directly observing the specimen immersed in a solution, needing no hardware modification.

**Fig. 4.**
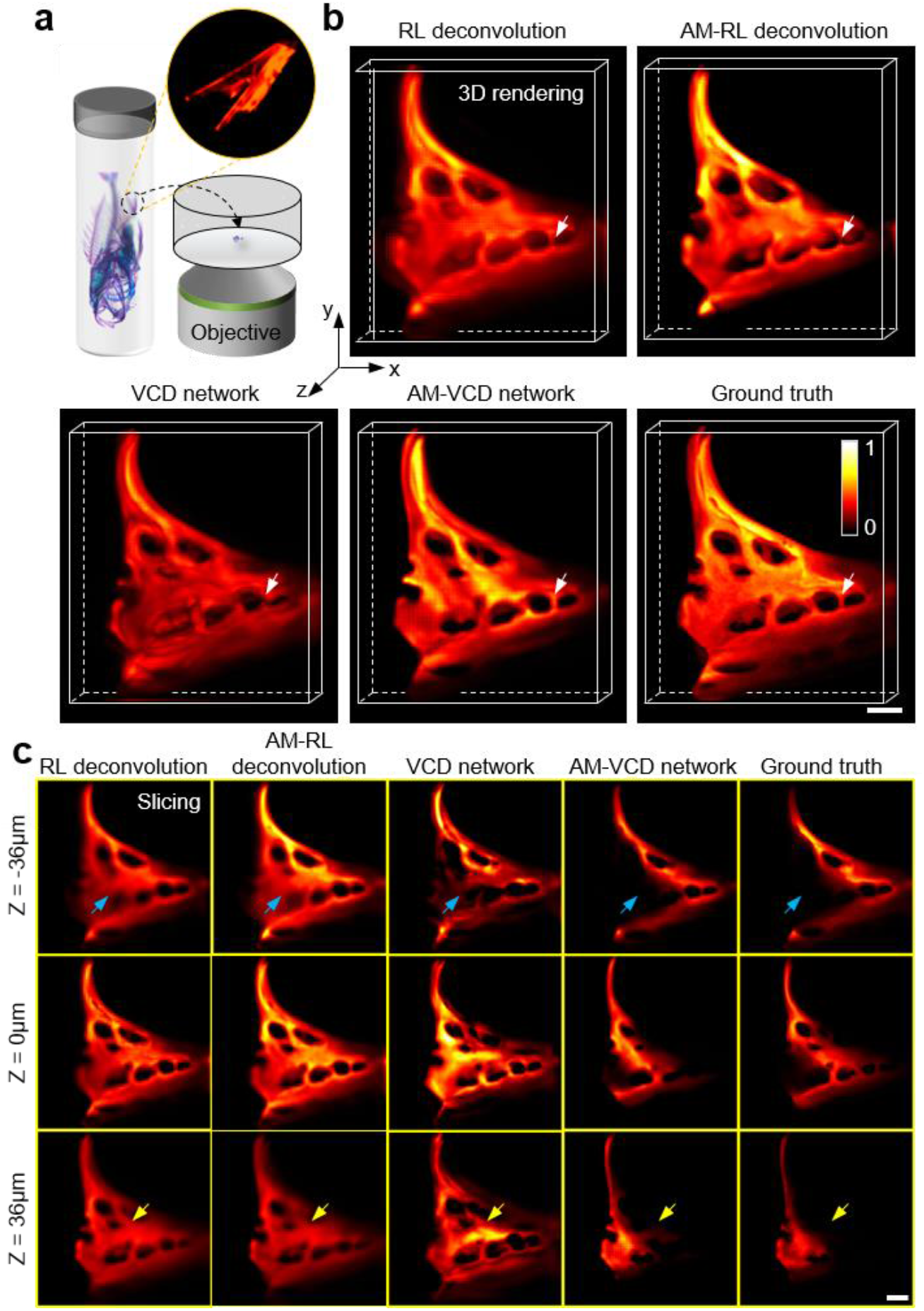
Imaging of fishbones of the glycerol-immersed largemouth bass sample by using an air-immersion objective lens (20×/0.5NA). **a.** A part of the largemouth bass specimen (hard bones dyed by Alizarin Red S) is immersed in the glycerol and placed in the glass disk for direct observation. **b.** 3D renderings of fishbones reconstructed by RL deconvolution, AM-RL deconvolution, VCD network, and AM-VCD network, respectively. The ground-truth 3D volume is acquired by confocal microscopy. Our AM-VCD network achieves the best performance with few artifacts, resolving the hole structures accurately **c.** Several slice images sectioned at three axial positions (z = - 36 μm, 0 μm, and 36 μm) of fishbones by RL deconvolution, AM-RL deconvolution, VCD network, AM-VCD network, and ground truth, respectively. Our AM-VCD network exhibits a strong optical sectioning ability consistent with the ground truth, as marked by blue and yellow arrows. Scale bar, 50 μm **b-c**.

### 3.4 AM-VCD enables resolution-uniform, artifact-suppress, and real-time 3D imaging of living samples

Our proposed AM-VCD network is very suitable for observing living samples that must be placed in the liquid, benefiting from its ability of directly imaging samples immersed in a solution and the inherent snapshot capability of LFM. In addition, our method can accurately reconstruct the 3D dynamics of samples in real-time and at micrometer resolution. Here we use experiments of dynamic imaging of zebrafish neutrophils (426×426×101 μm^3^ 3D volume and 6 frame rate) to demonstrate the superior capability of our method. As shown in Fig. 5a, after dropping the anesthetic into the living zebrafish larva placed in the glass disk, we immerse it with the 1% low-melting-temperature agarose solution. We use an air-immersion objective lens (20×/0.5NA) to continuously capture a series of LFM images to record the movement of neutrophils in blood vessels. Due to the different refractive indices between the agarose solution (refractive index = 1.335) and air (refractive index = 1) together with the thickness of the glass bottom (refractive index = 1.523), the spherical aberration is large, which is evaluated as −0.5λ. We use the AM-VCD network to reconstruct the 3D dynamics of zebrafish neutrophils from the series of LFM images. A reconstructed x-y MIP image at a specific moment (t = 59.00 s) is shown in Fig. 5a, where the bright-field image is captured and merged in as a background reference and different colors of neutrophils correspond to different axial depths. Our method can well resolve the point-structure of neutrophils in the 3D volume even under the large spherical aberration, showing the practicality and convenience of our method in real-time and 3D imaging of living samples. Fig. 5b shows the cell tracking results of reconstructed zebrafish neutrophils by the AM-VCD network, which correspond to the zoom-in area marked by the blue box in Fig. 5a. In Fig. 5c, some temporal sequences of the zoom-in area marked by the white box in Fig. 5a are also presented to show the dynamic imaging ability of our method (see Supplementary Video 1 for the whole moving process of zebrafish neutrophils in 3D). This experiment demonstrates that our AM-VCD network enables resolution-uniform, artifact-suppress, and real-time (high data acquisition speed and real-time reconstruction speed) 3D imaging of living samples in a solution, overcoming the limitation of the requirement of refractive index matching and improving the convenience of microscopic observation.

**Fig. 5.**
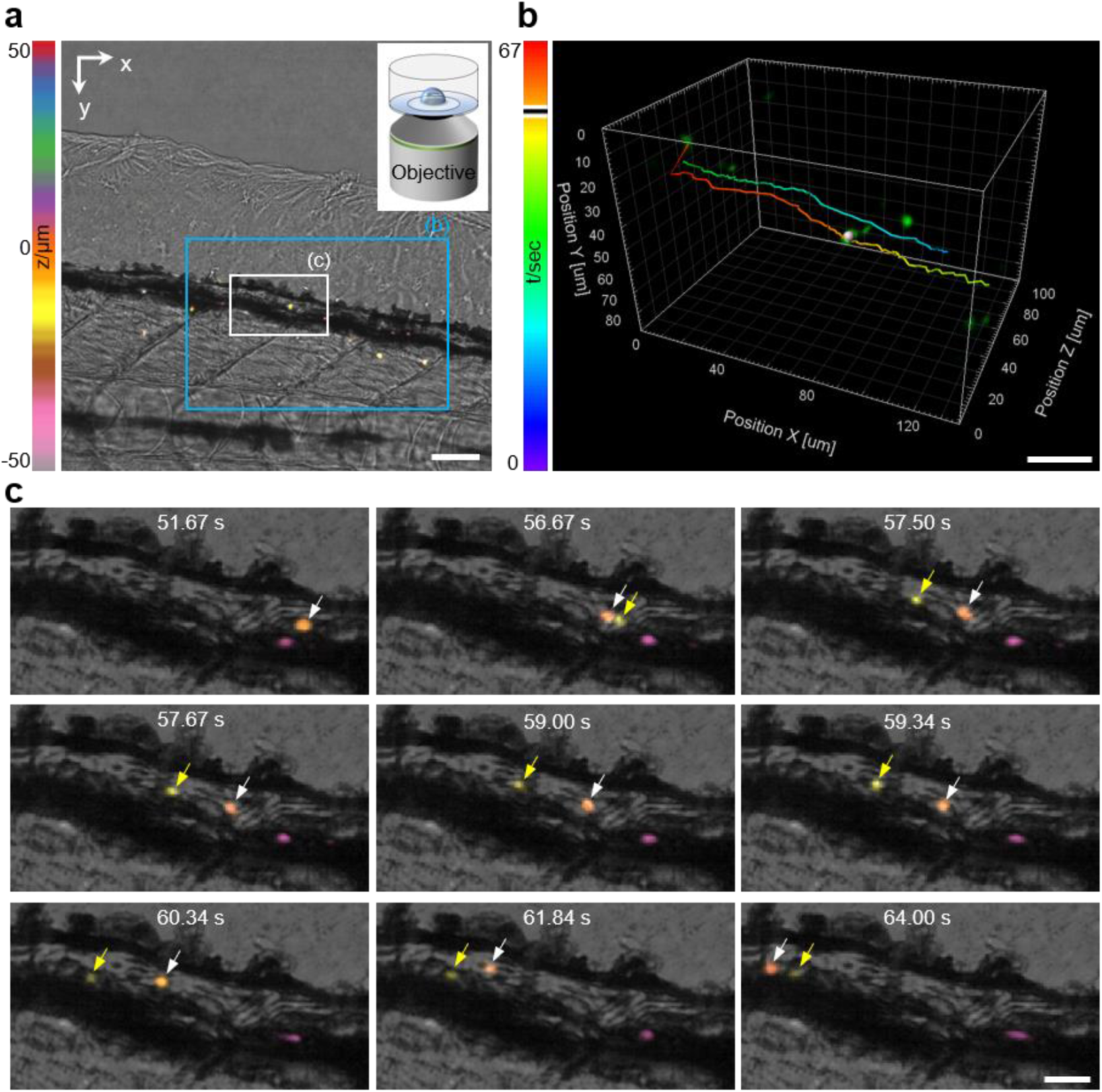
Dynamic 3D imaging of zebrafish neutrophils using an air-immersion objective lens (20×/0.5NA). **a.** The reconstructed lateral MIP at one certain moment (t = 59.00 s). A nonfluorescent bright-field image is captured as the reference and fused in the background. Up-right panel: the paralyzed zebrafish larva is immersed in 1% low-melting-temperature agarose solution and placed in a glass bottom dish. **b.** The cell tracking results of reconstructed zebrafish neutrophils by AM-VCD network, corresponding to the zoom-in area marked by the blue box in **a**. **c.** Time sequences of the zoom-in area marked by the white box in **a**. Scale bar, 50 μm **a**, 30 μm **b**, 10 μm **c**.

### 3.5 Data processing

For experiments using the20× objective, the 3D reconstruction comprises 101 axial slices with a voxel size of 0.33×0.33 × 1 μm^3^. For experiments using the 63× objective, the 3D reconstruction comprises 61 axial slices with a voxel size of 0.1 ×0.1 ×0.5 μm^3^.

In this work, we process data using MATLAB r2022b software in a 64-bit computer with Intel Core i9-9900X CPU @ 3.50GHz and 128GB memory and TensorFlow 1.0 in Nvidia GeForce RTX 2080 Ti graphic cards. The VCD and AM-VCD networks are trained on about 4000 pairs of image patches (176×176×101 pixel 3D volumes as ground truths and 176×176 pixel LFM images as the paired images), requiring 80 epochs and a time cost of approximately 6 hours on a single GPU. The reconstruction of a 671×671 pixel LFM image using the VCD and AM-VCD network both take about 70 ms for a single 3D volume, while RL deconvolution and AM-RL deconvolution (5 iterations) require approximately 300 seconds for reconstruction.

## 4 Discussion and Conclusion

In this work, we propose the AM-VCD network method to ease the performance reduction caused by the spherical aberration in LFM imaging. We model the aberration in the PSF generation of LFM and integrate it into the VCD deep learning structure. We further realize automatic aberration estimation and loss function optimization for the practical use of our method. Through the real experiments of fluorescent beads and numerical simulations under various aberration levels, we have proved that our method can achieve 3D volumetric reconstruction with artifact-free, real-time, resolution-uniform performance. We also prove that the proposed method can be used to directly and conveniently observe the largemouth bass sample immersed in the unmatched solution, resolving the complex structures of fishbones with high accuracy. Our method inherits the high acquisition speed of LFM and the real-time reconstruction speed of the VCD network. It only requires the conventional LFM system without no hardware modification and has the ability of overcoming great spherical aberration. Therefore, it is very suitable for long-term 3D dynamic observation of living samples placed in a solution, such as the imaging of living zebrafish without using exactly matched water-immersion objectives. In the future, we will introduce different types of aberration models into our method to expand its application scope and embed the aberration estimation step into the network structure to achieve more convenient and more precise aberration evolution.

## Supporting Information

Supporting Information is available from the Wiley Online Library or from the author.

## Acknowledgments

The work was supported by the National Science Foundation of China (NSFC) (Grant Nos. 62071219 and 62025108). X. Cao and Y. Zhou supervised this project. Y. Zhou and B. Xiong conceived and designed the experiments. Z. Jin and Q. Zhao prepared the samples. Z. Jin and Y. Zhou conducted the experiments. Z. Jin and B. Xiong realized the neural networks. Z. Jin analyzed and interpreted the experimental data. All authors discussed the results and contributed to the final manuscript.

## Data Availability Statement

The data and codes that support the figures and findings within this article are available from the corresponding authors upon reasonable request.

## Appendix A Sample Preparation

### Fluorescent beads

Fluorescent beads of 1-μm diameter (ABT-18-3-01, Bitoyscience) and 4-um diameter (ABT-18-3-04, Bitoyscience) were diluted 10000 times with distilled water and embedded on a glass bottom dish 15-mm diameter and 0.17-mm bottom thickness (NEST). The beads, which emit green fluorescence, were then imaged at room temperature.

### Largemouth bass bones imaging

For the preparation of largemouth bass specimens, we used the following reagents: distilled water, 95% ethanol, glycerol, 2% potassium hydroxide, hydrogen peroxide, and Alizarin Red S (emission wavelength is about 600 nm). We selected a largemouth bass with a length of 5 cm. To stiffen the fish, we first cleaned it with distilled water and then soaked it in 95% ethanol for 3-4 days. Next, we rehydrated the fish by placing it in distilled water for 24 hours. To transparentize the fish, we immersed it in a 2% potassium hydroxide solution for 24-48 hours. We then prepared a solution of Alizarin Red S by dissolving it in 95% ethanol and diluting it 10-fold with 1% potassium hydroxide solution. The fish was soaked in this solution for 48 hours to stain the bones. Finally, we preserved the fish specimen by immersing it in glycerol. To image the stained fish bones, we cut a small piece (about 2 mm×2 mm) from the fin and placed it in a glass bottom dish (15-mm diameter and 0.17-mm bottom thickness, NEST) filled with glycerol. We attached some green fluorescent beads to the surface of the fin slice. The emission peak of Alizarin Red in an aqueous medium is around 600 nm, which is also inside the wavelength range of the emission filter of the microscope, allowing us to simultaneously measure the experimental PSFs and image the fish bones. We used an air-immersion objective lens (20×/0.5NA) to capture images of the fish bones. To obtain high-resolution 3D data for network training, we imaged various parts of bones from the largemouth bass using an oil-immersion objective lens (40×/1.1NA Oil) on a confocal microscope (Zeiss, LSM 880).

### Dynamic imaging of zebrafish neutrophil

Transgenic zebrafish from the Tg(mpx:EGFP) line were utilized for neutrophil imaging. The embryos were incubated at 28.5°C until they reached 4 days post fertilization (dpf). To paralyze the larval zebrafish, the fish was briefly immersed in a solution of 1 mgml^−1^ alpha-bungarotoxin (Invitrogen). Once paralyzed, the larva was embedded in 1% low-melting-temperature agarose solution and placed in a glass bottom dish (15-mm diameter and 0.17-mm bottom thickness, NEST). In detail, we used a dropper to take about 1 ml of agar solution and dropped it into a glass bottom dish, then we used another dropper to move the zebrafish larva from the culture dish and fixed the zebrafish larva in the upper area of the droplet. We kept the specimen at room temperature for imaging.

