## Supplementary Material 1 for "Aberration modeling in deep learning for volumetric reconstruction of light-field microscopy"

**Supplementary Figures**

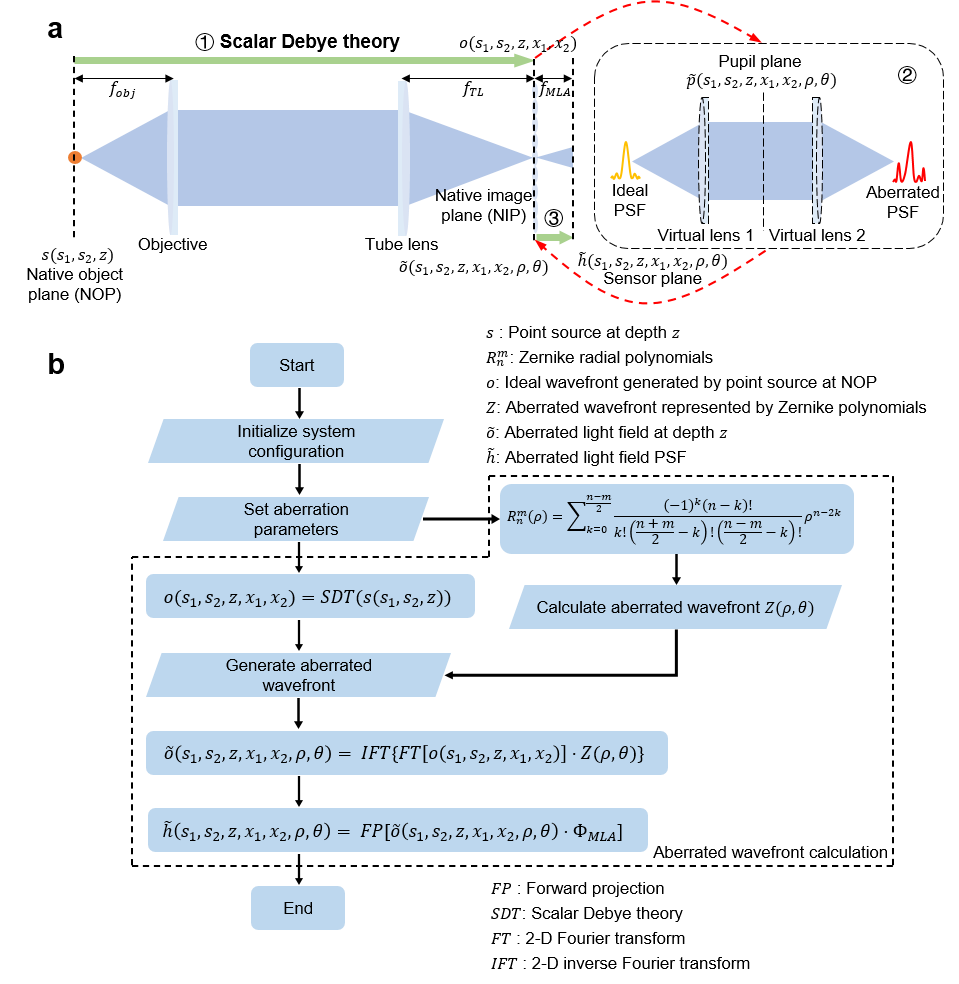

**Fig. S1. Aberration modeling in light-field microscopy (LFM).**

(**a**) In this forward projection (FP) model, the aberrated wavefront is generated by appending a virtual 4-f lens pair at the native image plane. Steps 1–3 illustrate this FP model, where we designate the object space coordinates as $(s_{1},s_{2},z)$, the pupil plane coordinates as $(\rho,\theta)$, and the sensor plane coordinates as $(x_{1},x_{2})$. **Step ①:** The diffraction-limited wavefront at the MLA plane (i.e. the ideal PSF before MLA), indicated as $o(s_{1},s_{2},z,x_{1},x_{2})$, is calculated using Scalar Debye Theory. **Step ②:** Appending a virtual 4-f lens pair to introduce the pupil plane into the optical model for phase modulation and the pupil function is $\tilde{p}(s_{1},s_{2},x_{1},x_{2},z,\rho,\theta)$. At the pupil plane, we use Zernike polynomials to describe the wavefront aberrations that existed in the anterior system as $Z(\rho,\theta)$. After pupil modulation, the aberrated wavefront at the MLA plane is $\tilde{o}(s_{1},s_{2},x_{1},x_{2},z,\rho,\theta)$. The calculation of pupil function is detailed in **Supplementary Note 1**. **Step ③:** The MLA at the native image plane (NIP) is modeled as a phase mask operating on this aberrated wavefront, which is then propagated to the sensor plane. The final wavefront at the sensor plane is $\tilde{h}(s_{1},s_{2},z,x_{1},x_{2},\rho,\theta)$, which indicates the aberrated PSF of the LFM system.

(**b**) Flowchart of aberration modeling in LFM. The abbreviations in use are described in the upper right and lower right corners.

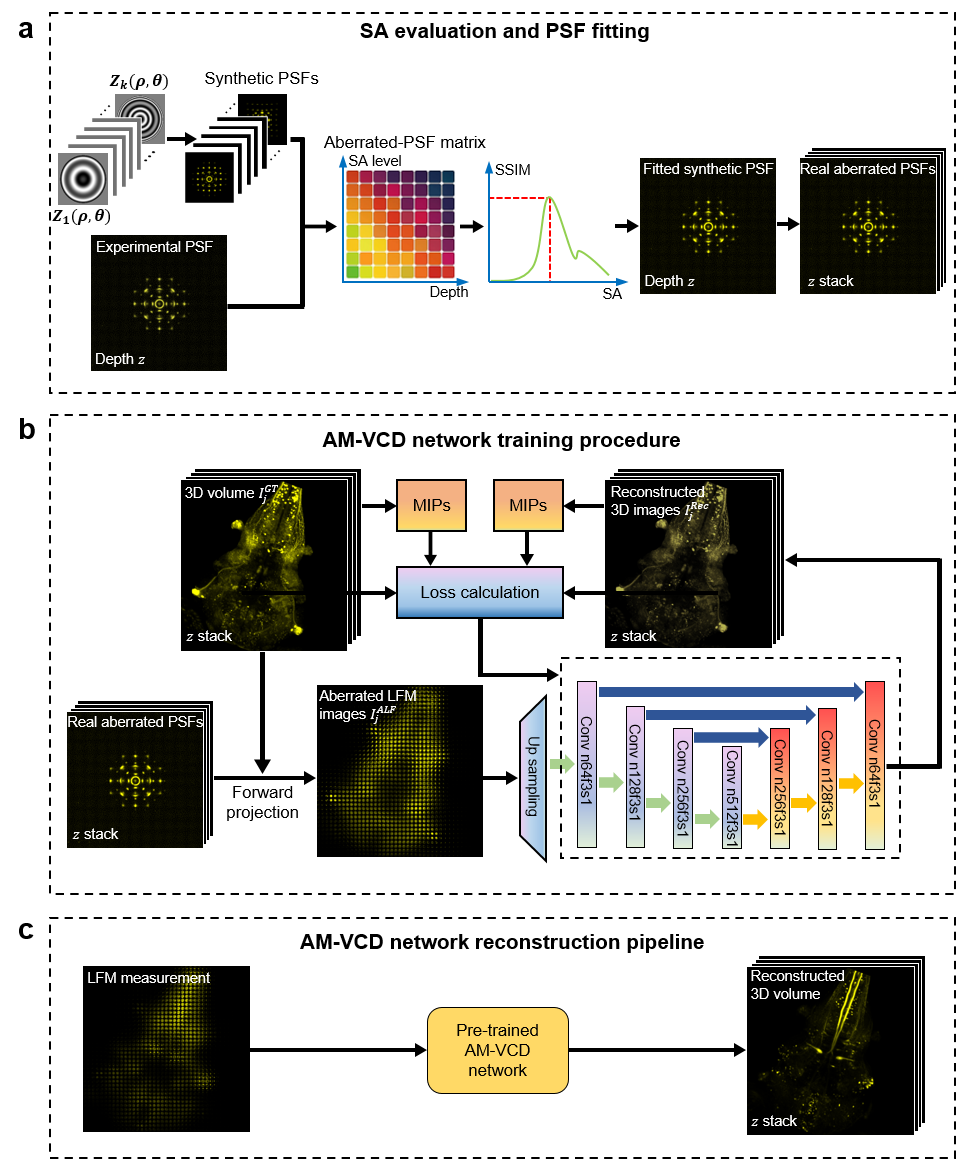

**Fig. S2. Training and reconstruction procedures of AM-VCD network.**

(**a**) Spherical aberration (SA) evaluation and real aberrated PSF stack generation procedure in LFM. To evaluate the SA level of a scene, (1) We first roughly estimate the depth range of the sample. We place fluorescent beads into the scene for observation or directly observe the bead-like structure inside the sample. These beads have less than or equal to the resolution limit of LFM. Since we find that the light-field PSF always has the minimum divergence at the focal plane regardless of the SA types and levels (see **Fig. S10** for details), we then adjust the objective lens axially until the observed bead appears in focus on the sensor to determine the focal plane (i.e., $z=0$). Next, we axially adjust the objective to a proper z depth for LFM imaging of the sample and record the current depth $z=z_{0}$. (2) We generate a series of aberrated wavefronts $Z_{i}\left( \rho,\theta\right) (i=1,2,\ldots,k)$ around the $z_{0}$ depth under different SA levels. By this, an aberrated-PSF matrix is calculated for PSF fitting, by comparing it with the experimentally-captured PSF image. We calculate the structural similarity indexes (SSIMs) of the experimental PSF and the synthetic PSFs from the aberrated-PSF matrix and select the synthetic PSF with the highest SSIM as the most fitted one. By this, the real SA level in the experiment is evaluated, which equals to that of the most fitted synthetic PSF. (3) Once the SA level is determined, a series of real aberrated PSFs with different z depths are generated based on the LFM parameters.

(**b**) The AM-VCD network training pipeline. We capture the ground-truth 3D data $I_{j}^{GT}$ by confocal or two-photon microscope. We use the ground-truth data to generate the corresponding aberrated LFM images $I_{j}^{ALF}$using the real aberrated PSFs with the evaluated SA level. The AM-VCD network is trained by iteratively minimizing the Mean Square Error (MSE) between the LFM reconstructed volumes and the ground truths. Maximum intensity projection (MIP) constraints, which indicate the weighted difference between the MIPs of the reconstructed volumes and the ground truths along the x-y, x-z, and y-z directions, are also added in the loss function. The loss function modification brings the artifact-reduction performance in reconstruction. Color blocks represent convolution layers with parameters n (channel number) and f (filter size). Blue arrows represent the concatenation operation.

(**c**) The newly captured LMF images can be fed into the pre-trained AM-VCD network for real-time 3D volume reconstruction.

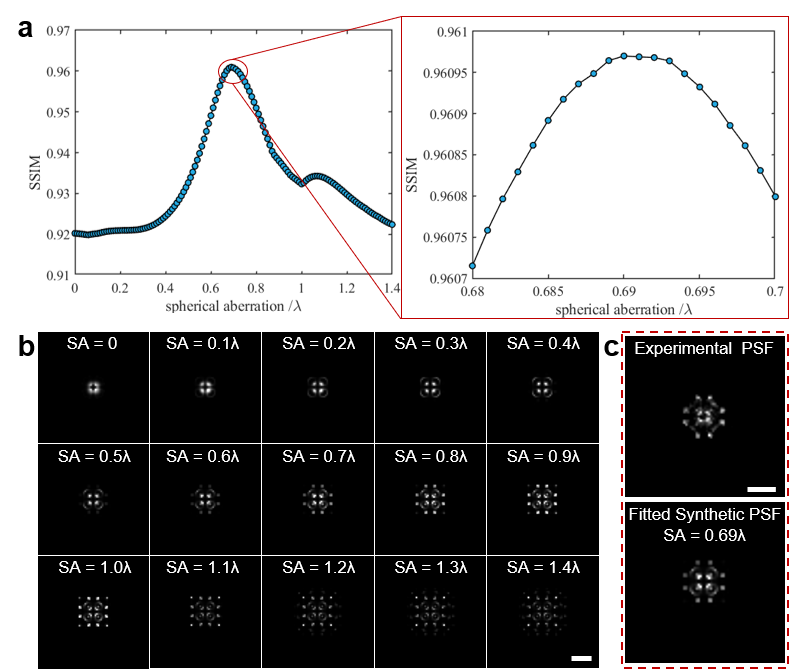

**Fig. S3. Aberration evaluation of an experimental scene.**

(**a**) The SSIM curve between the synthetic light-field PSFs and the real-captured experimental light-field PSF in the same depth (z = -10 μm here). Synthetic PSFs are generated with different SA levels ranging from 0 to 1λ (wavelength λ = 525 nm) with a step of 0.05λ. We compare the real-captured light-field PSF with the synthetic series to search for the most similar one, which we call the PSF fitting process. From the zoom-in panel of the curve, we can find that the highest SSIM value corresponds to the SA level of about 0.69λ, which indicates the real SA in the experiment.

(**b**) Some examples of synthetic light-field PSF images with different SA levels.

(**c**) Real-captured experimental PSF and its corresponding fitted synthetic PSF share very similar structures. Noting that the SA only needs to be estimated with a certain accuracy in real experiments, because the AM-VCD network has a robustness of reconstruction against minor estimated errors.

Scale bar, 10 μm (**b**), 5 μm (**c**).

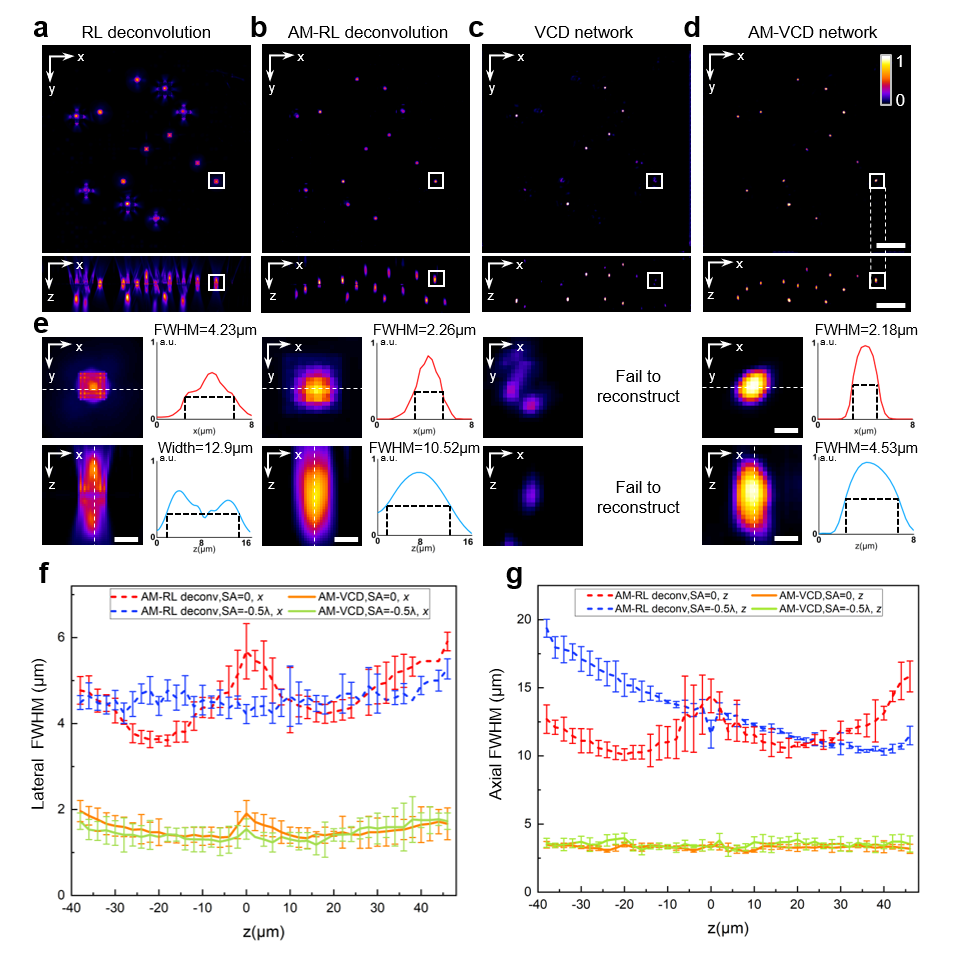

**Fig. S4 Performance and resolution evaluation of AM-VCD network (20×/0.5NA objective).**

We apply an air-immersion objective lens (20×/0.5NA) to image 4-μm diameter green fluorescent beads, which are placed in a glass slide. The SA level is estimated as -0.5λ, which is caused by the refractive index mismatch between the air (refractive index = 1) and the glass slide (refractive index = 1.523).

(**a**)-(**d**). Lateral and axial MIPs of real-captured fluorescent beads reconstructed by RL deconvolution, AM-RL deconvolution, VCD network, and AM-VCD network, respectively.

(**e**) Zoom-in areas marked by white boxes in (**a**)**-**(**d**). The profiles along dashed lines are plotted and full width at half maximum (FWHM) values are calculated in both lateral and axial directions (a.u., arbitrary unit).

(**f**)-(**g**). The average lateral FWHM and average axial FWHM of simulated 3D-distributed beads under different SA levels (0 and -0.5λ) are reconstructed using both AM-RL deconvolution (dashed lines) and AM-VCD network (solid lines) with corresponding aberrated PSFs.

In both experiments and simulations, methods that incorporate aberration modeling, such as AM-RL deconvolution and AM-VCD network, lead to improved reconstruction resolution. AM-VCD network can further enhance the spatial resolution in both the lateral and axial directions, and can effectively reduce reconstruction errors by the original VCD network.

Scale bar, 10 μm (**a**)-(**d**).

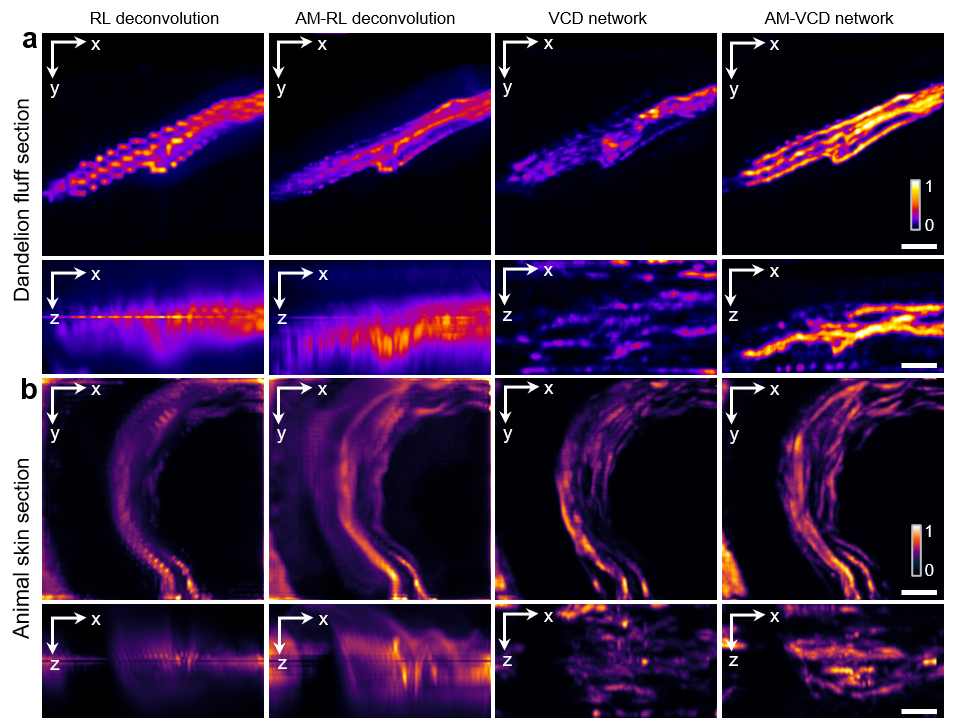

**Fig. S5 More experimental results by different reconstruction methods in imaging biological slices (63×/1.25NA objective).**

Here we apply a 63×/1.25NA oil-immersion objective lens to image biological samples immersed in water, which have large SAs. To quantitatively measure the SA, we drop some fluorescent beads on the cover slip and capture a snapshot LFM image. The evaluated aberration level is 0.8λ, which is then applied in both AM-RL deconvolution and AM-VCD network reconstruction.

(**a**) Lateral and axial MIPs of unstained dandelion villi slice.

(**b**) Lateral and axial MIPs of an H&E-stained animal skin slice.

From the results of MIPs, we can see that the AM-VCD network achieves the best imaging performance with the least reconstruction artifacts and the highest spatial resolution in both lateral and axial directions.

Scale bar, 10 μm.

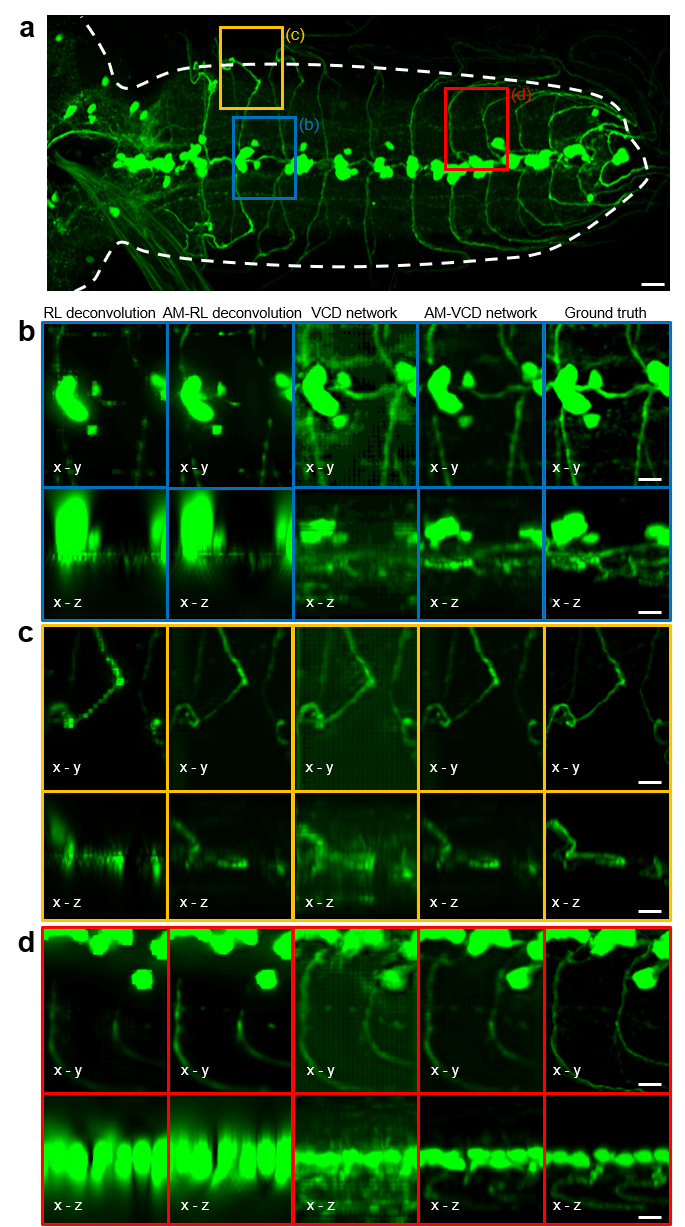

**Fig. S6 Simulation comparison of reconstruction performance by different reconstruction methods in imaging Drosophila larvae.**

(**a**) x-y MIP image of a full-FOV 3D volume of drosophila larvae (Tdc2-LexA) from the ground-truth dataset, which is captured by a two-photon microscope.

(**b**)-(**d**) Reconstructed MIP images of the synthetic LFM images under an SA level of 0.3λ by using RL deconvolution, AM-RL deconvolution, VCD network, and our AM-VCD network. These reconstructed sub-region images correspond to the areas in (**a**) marked by different color boxes. The results show that the AM-VCD network can recover the high-fidelity 3D distributions of drosophila larvae with complex structures. It provides higher spatial resolution than deconvolution methods, especially in the axial direction. It can also effectively suppress the periodic artifacts by the VCD network reconstruction. The parameters are set the same as those in real experiments using the 20×/0.5NA objective lens**.**

Scale bar, 20 μm (**a**), 10 μm (**b**)-(**d**).

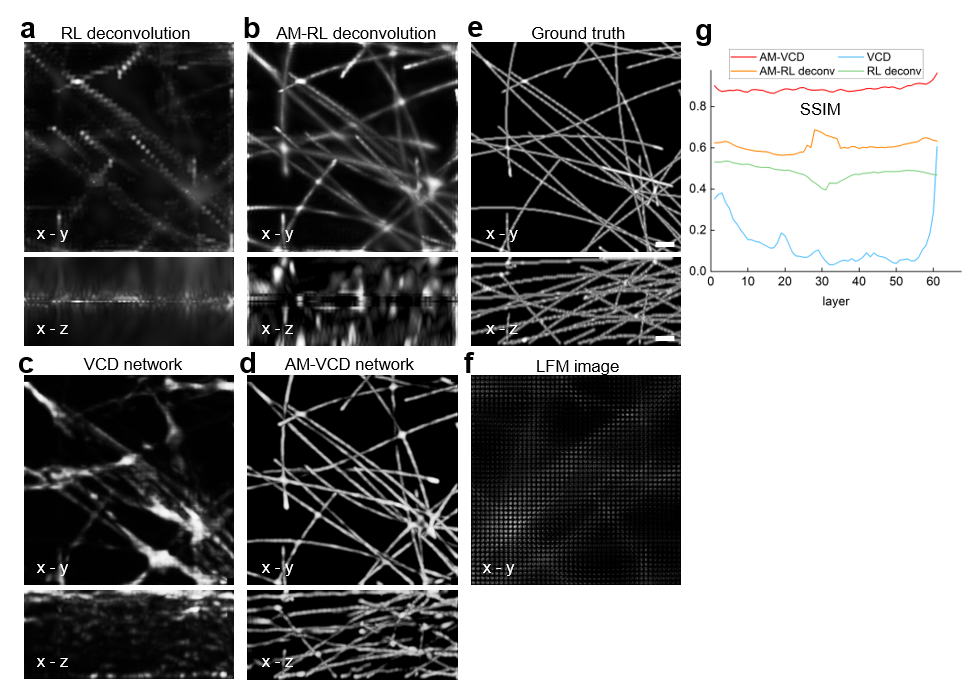

**Fig. S7. Simulation verification of the robust reconstruction ability of AM-VCD network under an extremely large SA level.**

Here we use the simulation of tubulins data^1^ to demonstrate the reconstruction ability of our AM-VCD network under an extremely large SA level (1.6λ). Other parameters are set the same as those in real experiments using the 63×/1.25NA objective lens.

(**a**)-(**d**) Reconstructed MIP images of tubulins data from a snapshot synthetic LFM image with a SA level of 1.6λ by using different methods, such as RL deconvolution, AM-RL deconvolution, VCD network, and AM-VCD network.

(**e**) The MIP image of the corresponding 3D volume from the ground-truth tubulins dataset^1^.

(**f**) Synthetic aberrated LFM image generated from the ground-truth 3D volume.

(**g**) The SSIM index curves exhibit the reconstructed results in different axial layers by using different methods, referring to the ground truth.

The results of MIPs and SSIMs show that the AM-VCD can reconstruct an acceptable result even under this extremely large SA, while other methods can hardly obtain a reliable performance.

Scale bar, 50 μm (**a**)-(**f**).

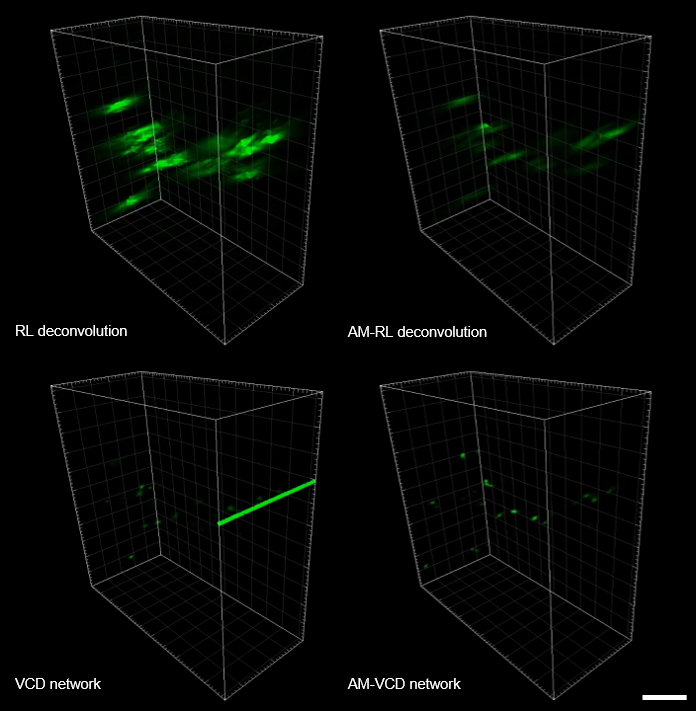

**Fig. S8. Reconstruction comparison of zebrafish neutrophil cells by using RL deconvolution, AM-RL deconvolution, VCD network, and AM-VCD network (20×/0.5NA objective).**

Here we show the reconstructed volumes of zebrafish neutrophils at one certain moment (t = 13.193 s) by using different methods. The results show that RL deconvolution and VCD network both fail to reconstruct the dynamics of zebrafish neutrophils. The result of RL deconvolution has poor lateral and axial resolutions, while the result of VCD has heavy errors and artifacts (see the green thick line). The AM-RL deconvolution can correctly resolve the zebrafish neutrophils in the 3D volume but still has a low axial resolution compared to the proposed AM-VCD network (see **Supplementary Video 1** for the full dynamic process).

Scale bar, 40 μm.

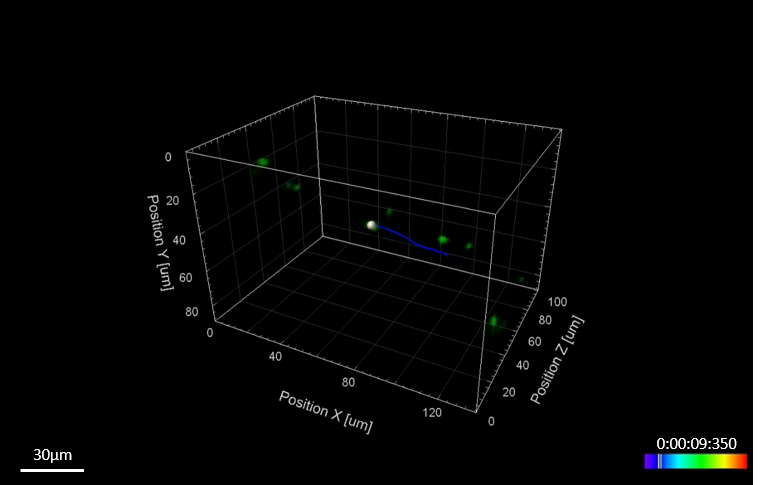

**Fig. S9. Zebrafish neutrophil cell tracking result by AM-VCD network (20×/0.5NA objective).**

Here we show the cell tracking result of reconstructed Zebrafish neutrophil by AM-VCD network at one certain moment (t = 9.350 s). The total tracing duration is 2 seconds, indicated by the blue line. We apply the software (Imaris 9.0.1 software) to detect the moving cells automatically and assign time-stamped color codes to each cell trace. The tracked cells are then displayed with white spheres overlaid on top (see **Supplementary Video 1** for the full tracing process).

Scale bar, 30 μm.

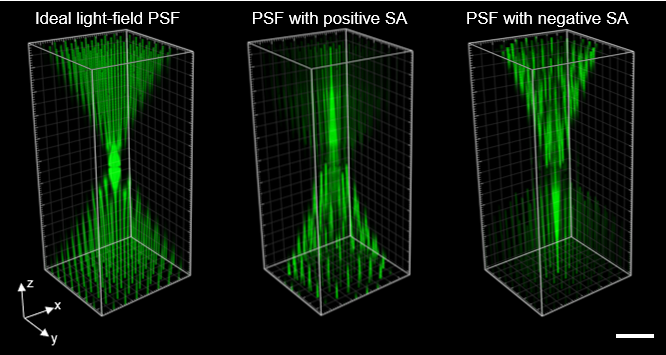

**Fig. S10. Sampling pattern comparison between ideal light-field PSF, light-field PSF with a positive SA (SA = 0.5λ), and light-field PSF with a negative SA (SA = -0.5λ).**

The appearance of SA changes the sampling pattern of light-field PSF. For an ideal LFM system, the sampling pattern is relatively uniform and symmetric on both sides of the focal plane. For the system with a positive SA, the sampling becomes denser on one side (above the focal plane, as shown in the figure) and sparser on another side (below the focal plane, as shown in the figure), which is the opposite for the system with a negative SA. This phenomenon of sampling change will cause the resolution change along the imaging depth, which can be found in the reconstructed results by AM-RL deconvolution in Fig. 2g and Fig. S4g. Specifically, the axial positions with the highest resolution and lowest resolution are quite different from the original LFM system. However, we can also observe that regardless of the SA types and levels, the light-field PSF always has the minimum divergence at the focal plane.

Scale bar, 10 μm.

**Supplementary Note 1: Calculation of aberrated pupil function** $\tilde{p}\left( \rho,\theta\right)$

First, we give the generation process of the wavefront function $Z(\rho,\theta$), which has been mentioned in Fig. S1. The wavefront function $Z(\rho,\theta$) in this work is introduced by aberrations and generally can be calculated by a weighted combination of normalized Zernike polynomials^2^.

Since the optical system in this work is assumed to be circular in shape, we use polar coordinates for formula representation. The definition of the Zernike polynomials^2^ is as follows

$$\begin{aligned} \left\{ \begin{aligned} Z_{n}^{m}\left( \rho,\theta\right)={\sqrt{2(n+1)} R}_{n}^{m}\left( \rho\right)\cos\left( m\theta\right), even Zernike polynomials \\ Z_{n}^{-m}\left( \rho,\theta\right)={\sqrt{2(n+1)} R}_{n}^{m}\left( \rho\right)\sin\left( m\theta\right), odd Zernike polynomials \end{aligned} \right.\#(S1) \end{aligned}$$

where $n\geq m\geq0, n,m\in\mathbf{N}$. $\rho(0\leq\rho\leq1)$ represents the radial distance and $\theta$ is the azimuthal angle.$R_{n}^{m}$ is the radial polynomials and can be expressed as

$$\begin{aligned} R_{n}^{m}\left( \rho\right)=\sum_{k=0}^{\frac{n-m}{2}} \frac{\left( -1 \right)^{k}\left( n-k \right)!}{k!\left( \frac{n+m}{2}-k \right)!\left( \frac{n-m}{2}-k \right)!}\rho^{n-2k}.\#\left( S2 \right) \end{aligned}$$

Since we only consider the primary spherical aberration in this work, the wavefront function can be simplified with only the normalized primary spherical item, which can be written as

$$\begin{aligned} {Z\left( \rho,\theta\right)=Z}_{4}^{0}\left( \rho,\theta\right)=\sqrt{5}\left( 6\rho^{4}-6\rho^{2}+1 \right).\#\left( S3 \right) \end{aligned}$$

The aberrated pupil function $\tilde{p}$is the joint operation of the wavefront function $Z(\rho,\theta$) and a circular cut-off function (limited by the NA of the objective lens), which can be expressed as

$$\begin{aligned} \tilde{p}\left( \rho,\theta\right)={e^{-ikZ\left( \rho,\theta\right)}|}_{\rho\leq r_{xp}}.\#\left( S4 \right) \end{aligned}$$

We can further rewrite Equation (S4) in the rectangular coordinate system for easy use as follows

$$\begin{aligned} \tilde{p}\left( x,y \right)=\mathrm{circ}\left( \frac{\sqrt{x^{2}+y^{2}}}{r_{xp}} \right)e^{-ikZ\left( x,y \right)},\#\left( S5 \right) \end{aligned}$$

where$rect(\cdot)$ indicates the rectangle function, $x=\rho\cos\theta$ and $y=\rho\sin\theta$ are the normalized coordinates at the exit pupil plane. $k=\frac{2\pi}{\lambda}$ is the wave number and$r_{xp}$ is the radius of the exit pupil, where $\lambda$ is the wavelength of light.
